# The CRISPR spacer space is dominated by sequences from the species-specific mobilome

**DOI:** 10.1101/137356

**Authors:** Sergey A. Shmakov, Vassilii Sitnik, Kira S. Makarova, Yuri I. Wolf, Konstantin V. Severinov, Eugene V. Koonin

## Abstract

The CRISPR-Cas is the prokaryotic adaptive immunity system that stores memory of past encounters with foreign DNA in spacers that are inserted between direct repeats in CRISPR arrays ^1,2^. Only for a small fraction of the spacers, homologous sequences, termed protospacers, are detectable in viral, plasmid or microbial genomes ^3,4^. The rest of the spacers remain the CRISPR “dark matter”. We performed a comprehensive analysis of the spacers from all CRISPR-*cas* loci identified in bacterial and archaeal genomes, and found that, depending on the CRISPR-Cas subtype and the prokaryotic phylum, protospacers were detectable for 1 to about 19% of the spacers (∼7% global average). Among the detected protospacers, the majority, typically, 80 to 90%, originate from viral genomes, and among the rest, the most common source are genes integrated in microbial chromosomes but involved in plasmid conjugation or replication. Thus, almost all spacers with identifiable protospacers target mobile genetic elements (MGE). The GC-content, as well as dinucleotide and tetranucleotide compositions, of microbial genomes, their spacer complements, and the cognate viral genomes show a nearly perfect correlation and are almost identical. Given the near absence of self-targeting spacers, these findings are best compatible with the possibility that the spacers, including the dark matter, are derived almost completely from the species-specific microbial mobilomes.

---

Driven by the overwhelming success of the Cas9 and later Cpf1 endonucleases as the new generation of genome editing tools, comparative genomics, structures, biochemical activities and biological functions of CRISPR (Clustered Regularly Interspaced Palindromic Repeats)-Cas (CRISPR-associated proteins) systems have been recently explored in unprecedented detail ^1,2,5,6^. The CRISPR-Cas are adaptive (acquired) immune systems of archaea and bacteria that store memory of past encounters with foreign DNA in unique spacer sequences that are excised from viral and plasmid genomes by the Cas adaptation machinery, or alternatively, reverse transcribed from foreign RNA and inserted into CRISPR arrays ^7,8^. Transcripts of the spacers, together with portions of the surrounding repeats, are employed by Cas effector complexes as guide CRISPR (cr)RNAs to recognize the cognate sequences (protospacers) in the foreign genomes upon subsequent encounters, directing Cas nucleases to their cleavage sites ^9,10^ and limiting bacteriophage infection and horizontal gene transfer.^REF^

One of the burning open questions in the CRISPR area is the origin of the bulk of the spacers. For a small fraction of the spacers, protospacers have been reported, often in viral and plasmid genomes, but the overwhelming majority of the spacers remain without a match ^3,4,11^-^15^. In order to get insight into the origin of this “dark matter”, we performed comprehensive searches of the current genomic and metagenomic sequence databases using all identifiable spacer sequences from complete bacterial and archaeal genomes as queries. To this end, a computational pipeline was developed that identified all CRISPR arrays from complete and partial bacterial and archaeal genomes, extracted the spacers and used them as queries to search the viral and prokaryotic subsets of the Non-redundant nucleotide database at the NCBI (NIH, Bethesda) for protospacers under stringent criteria for homology detection (Supplementary Figure 1 and Supplementary text 1; see Methods for details).

These searches yielded 2,981 spacer matches (protospacers) in viral sequences and 23,385 matches in prokaryotic sequences. We then examined the provenance of the detected protospacers across the diversity of the CRISPR-Cas systems and the prokaryotic phyla. In a general agreement with previous analyses that, however, have been performed on much smaller genomic data sets, protospacers were identified for ∼7% of the spacers, with the fractions for different CRISPR-Cas subtypes ranging from 1 to 19% (Table 1). The fraction of detected protospacers was typically higher for type I and II CRISPR-Cas systems, in which it spans the entire range, compared to type III, where this fraction was uniformly low, at 1 to 2% (Table 1).

**Table 1.**
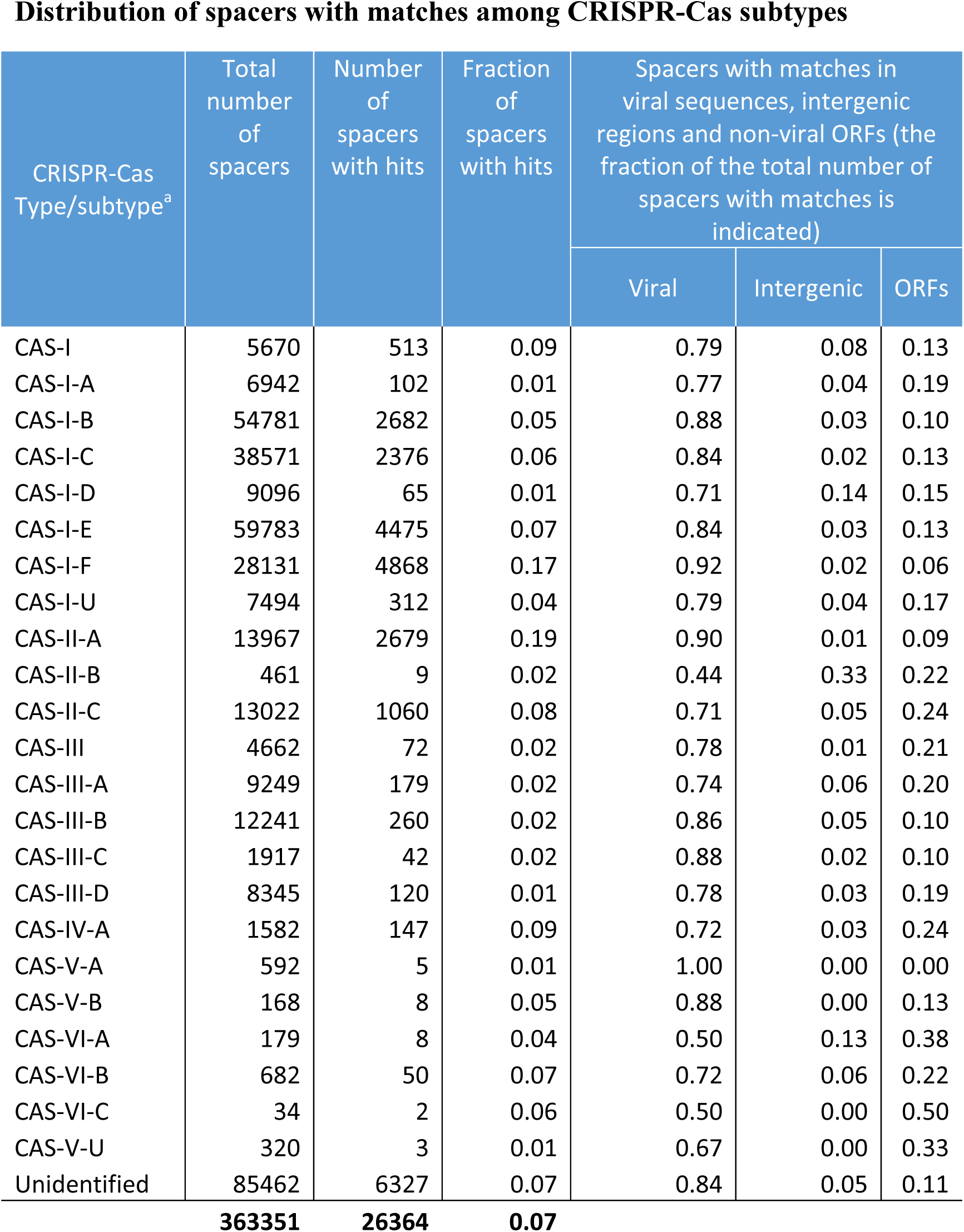
Distribution of spacers with matches among CRISPR-Cas subtypes. Identification and classification of the CRISPR-Cas systems were as previously described ^16,42^; CAS-I, CAS-III denote loci that could be assigned to types I and III, respectively, but not to a specific subtype; Unidentified are orphan CRISPR arrays and incomplete CRISPR-*cas* loci.

A similar range was detected for the fraction of spacers with matches across the bacterial and archaeal phyla (Table 2) but substantial deviations from the global average of ∼7% in several phyla are notable. Thus, anomalously high fractions of spacers with matches were detected in *Spirochaetia, Fusobacteria and γ-Proteobacteria.* In a sharp contrast, the CRISPR arrays in archaea, especially hyperthermophiles, had low fraction of matching spacers, with none at all detected in *Thermococci* and *Thermoplasmata*; furthermore, the only phylum of hyperthermophilic bacteria, for which a large number of CRISPR arrays was identified, also had only 1% of matching spacers (Table 2). A multiple regression analysis shows that both the assignment to a CRISPR subtype and classification into an archaeal or bacterial phylum make substantial and largely independent contributions to the variation of the fraction of spacers with detectable matches; jointly, the two factors explain about 75% of the variance of that fraction (see Supplementary text 1). The paucity of spacer matches in hyperthermophiles is puzzling because all these organisms possess CRISPR-*cas* loci (as opposed to only a minority among mesophiles) ^16^, with the implication that CRISPR activity is essential for the survival of these organisms. The lack of recognizable spacers could be due to under-sampling of the respective virome and/or to preferential utilization of partially matching spacers by the CRISPR-Cas systems of thermophiles. Generally, the aspects of the biology of different groups of prokaryotes that might determine the activity of the CRISPR-Cas systems, and hence the fraction of spacers with matches, remain to be explored.

**Table 2.** Distribution of spacers with matches among bacterial and archaeal phyla.

| Phylum | Total Number of spacers<sup>a</sup> | Number of spacers With matches<sup>a</sup> | Fraction of spacers with matches | Spacers with matches in viral sequences, intergenic regions and non-viral ORFs (the fraction of the total number of spacers with matches is indicated) |  |  |
| --- | --- | --- | --- | --- | --- | --- |
|  |  |  |  | Viral | Intergenic | ORFs |
| Actinobacteria | 54875 | 3614 | 0.07 | 0.76 | 0.05 | 0.19 |
| Alphaproteobacteria | 8135 | 120 | 0.01 | 0.69 | 0.07 | 0.24 |
| Bacteroidetes/Chlorobi group | 18611 | 840 | 0.05 | 0.78 | 0.03 | 0.19 |
| Betaproteobacteria | 14013 | 908 | 0.06 | 0.69 | 0.14 | 0.16 |
| Chloroflexi | 6523 | 30 | 0.00 | 0.77 | 0.03 | 0.20 |
| Crenarchaeota | 11212 | 119 | 0.01 | 0.90 | 0.02 | 0.08 |
| Cyanobacteria/Melainabacteria group | 20295 | 126 | 0.01 | 0.75 | 0.04 | 0.21 |
| Deinococcus-Thermus | 4057 | 85 | 0.02 | 0.75 | 0.04 | 0.21 |
| delta/epsilon subdivisions | 13588 | 378 | 0.03 | 0.60 | 0.06 | 0.34 |
| Firmicutes | 93332 | 7643 | 0.08 | 0.90 | 0.02 | 0.08 |
| Fusobacteriia | 3427 | 629 | 0.18 | 0.92 | 0.01 | 0.06 |
| Gammaproteobacteria | 67202 | 10238 | 0.15 | 0.91 | 0.03 | 0.06 |
| Halobacteria | 5121 | 74 | 0.01 | 0.55 | 0.08 | 0.36 |
| Methanobacteria | 2218 | 47 | 0.02 | 0.70 | 0.04 | 0.26 |
| Methanococci | 1639 | 6 | 0.00 | 0.50 | 0.00 | 0.50 |
| Methanomicrobia | 10399 | 141 | 0.01 | 0.91 | 0.02 | 0.06 |
| Nitrospira | 1088 | 13 | 0.01 | 0.85 | 0.00 | 0.15 |
| Planctomycetes | 1650 | 14 | 0.01 | 0.79 | 0.14 | 0.07 |
| Spirochaetia | 5114 | 1173 | 0.23 | 0.73 | 0.04 | 0.24 |
| Synergistia | 1702 | 22 | 0.01 | 0.64 | 0.00 | 0.36 |
| Tenericutes | 1050 | 26 | 0.02 | 0.73 | 0.04 | 0.23 |
| Thermococci | 3210 | 16 | 0.00 | 0.31 | 0.00 | 0.69 |
| Thermoplasmata | 1270 | 6 | 0.00 | 0.17 | 0.17 | 0.67 |
| Thermotogae | 3731 | 31 | 0.01 | 0.94 | 0.00 | 0.06 |
| unclassified Bacteria (miscellaneous) | 2814 | 6 | 0.00 | 0.67 | 0.00 | 0.33 |
|  | **356276** | **26305** | **0.07** |  |  |  |

<sup>a</sup>Only phyla with >1,000 unique spacers were included, hence slightly lower total number of spacers than in Table 1.

The CRISPR-Cas spacers have been demonstrated to insert in a polarized fashion, mostly in the beginning of arrays, adjacent to the leader sequence (although in some case, internal insertion has been observed as well), resulting in unidirectional growth of the array that, however, subsequently contracts via loss of distal spacers ^17,18^. Indeed, a notable excess of spacers with matches was observed near the ends of the arrays, with a sharp decline downstream (Figure 1A,B), indicating that a large fraction of recently acquired spacers originate from sequences available in current databases.

**Figure 1.**
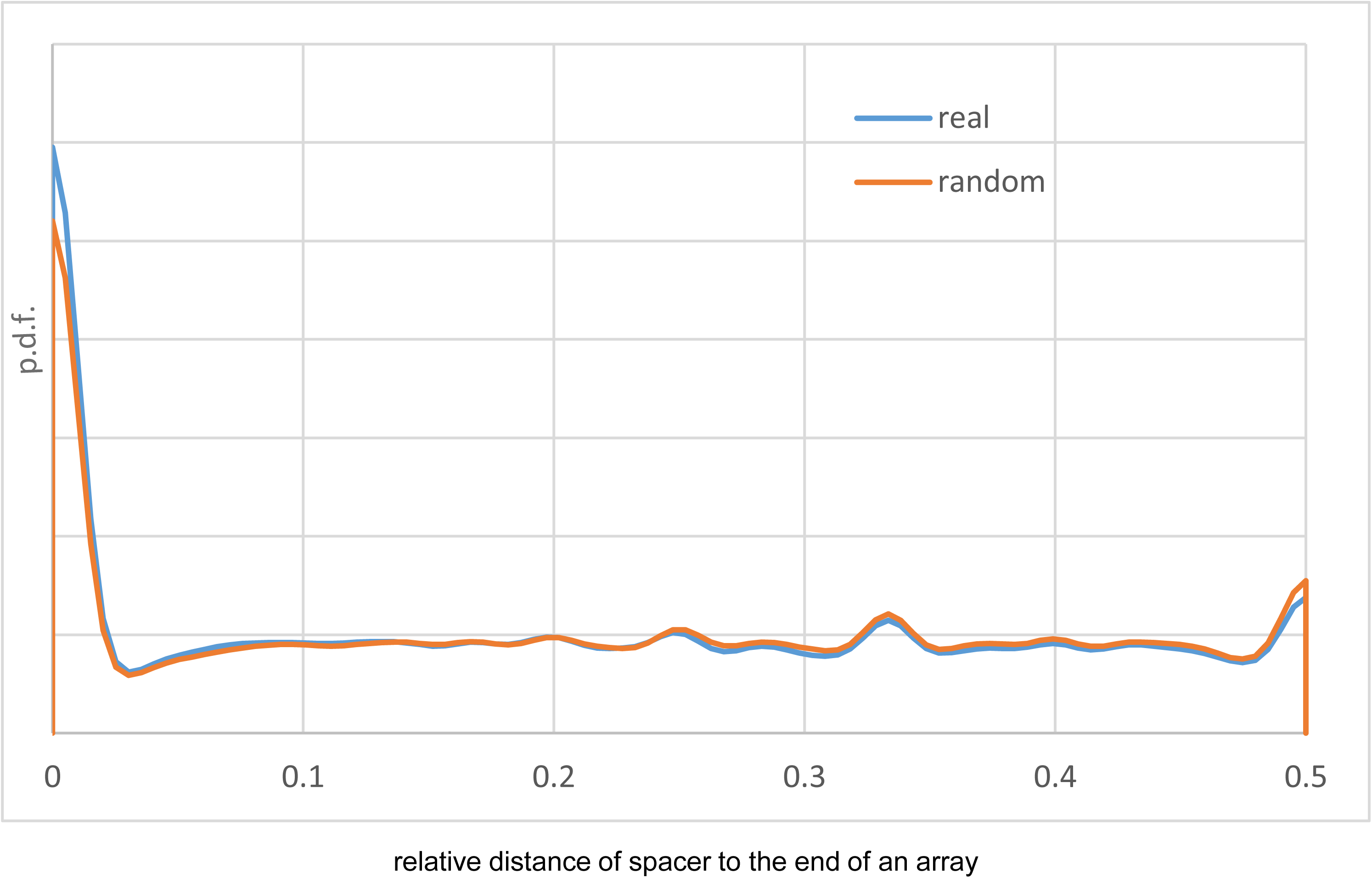
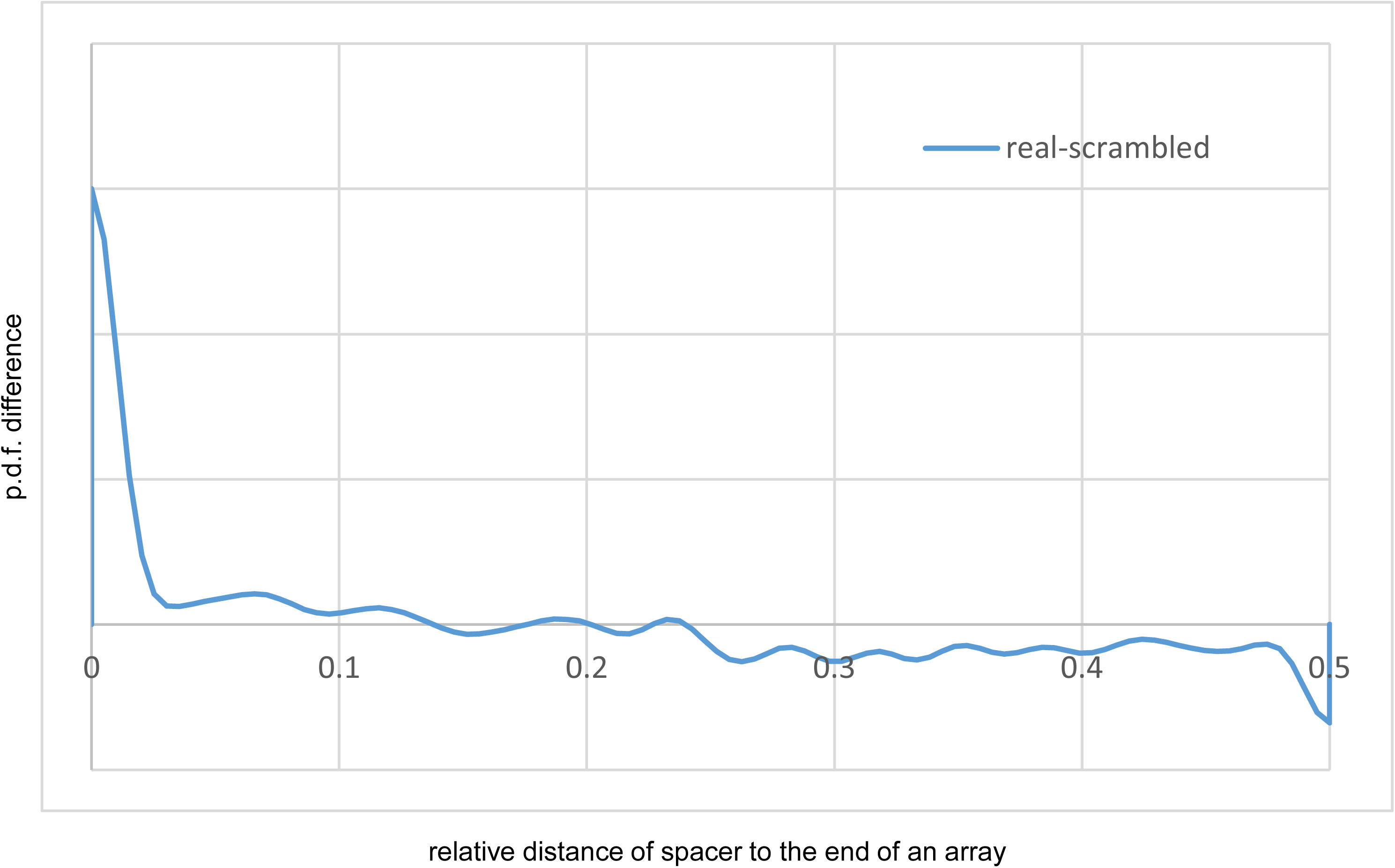
Distribution of the spacers with matches along the CRISPR arrays. (A) Probability density functions for the spacers with matches (real) and for the same spacers placed randomly onto the array 100 times (random). (B) Probability density function of the difference between the number of spacers with matches and randomly placed spacers along the array. Given the difficulty of polarizing CRISPR arrays automatically and under the assumption that new spacers are incorporated at the leader end but not at the distal end of arrays, the results are shown from the end to the middle of the arrays.

In most subtypes of CRISPR-Cas from most bacterial and archaeal phyla, 70 to 90% of the protospacers originated from virus or provirus sequences (proviruses were consistently identified with two independent approaches; see Supplementary figure 2 and Methods for details) (Tables 1 and 2), in agreement with the common notion that CRISPR-Cas is primarily engaged in antiviral defense. Notably, subsets of virus-specific spacers are shared between different species and even genera of bacteria (e.g. *Staphylococcus*-*Streptococcus* and *Escherichia*-*Cronobacter*), which yields a host-virus network that includes several large connected components (Supplementary Figure 3, Supplementary data set 1). Analysis of the provenance of the non-viral protospacers showed a clear preponderance of sequences from gene families implicated in conjugal transfer and replication of plasmids, such as type IV secretion systems ^19^ (Figure 2 and Supplementary data set 2). Notably, several protospacers also originated from *cas* genes, particularly *cas3* (Figure 2 and Supplementary Table 1), recapitulating the recent finding of *cas*-matching protospacers in orphan CRISPR arrays ^20^. Of the remaining genes containing protospacers, many are unannotated, which is typically caused by low sequence conservation, and potentially could originate from viruses or plasmids as well. A small fraction of spacer matches map to genomic regions annotated as intergenic (Tables 1 and 2) but manual examination of such cases led to identification of putative protein-coding genes that apparently have been missed by genome annotation (Supplementary text 2). Complete reannotation of the available prokaryotic genomes is a demanding project outside the scope of this work but, with this caveat, only a small fraction of the detected protospacers could be traced to sequences demonstrably not originating from viruses or other mobile elements. Previous analyses of CRISPR arrays from individual bacterial and archaeal genomes have reported widely different fractions of self-matching spacers 1,21. Our current, comprehensive analysis indicates that the overwhelming majority of the spacers that persist long enough to be detected are derived from viruses and other mobile elements (collectively, known as the mobilome ^22^), apparently indicating strong selection against self-targeting spacers.

**Figure 2.**
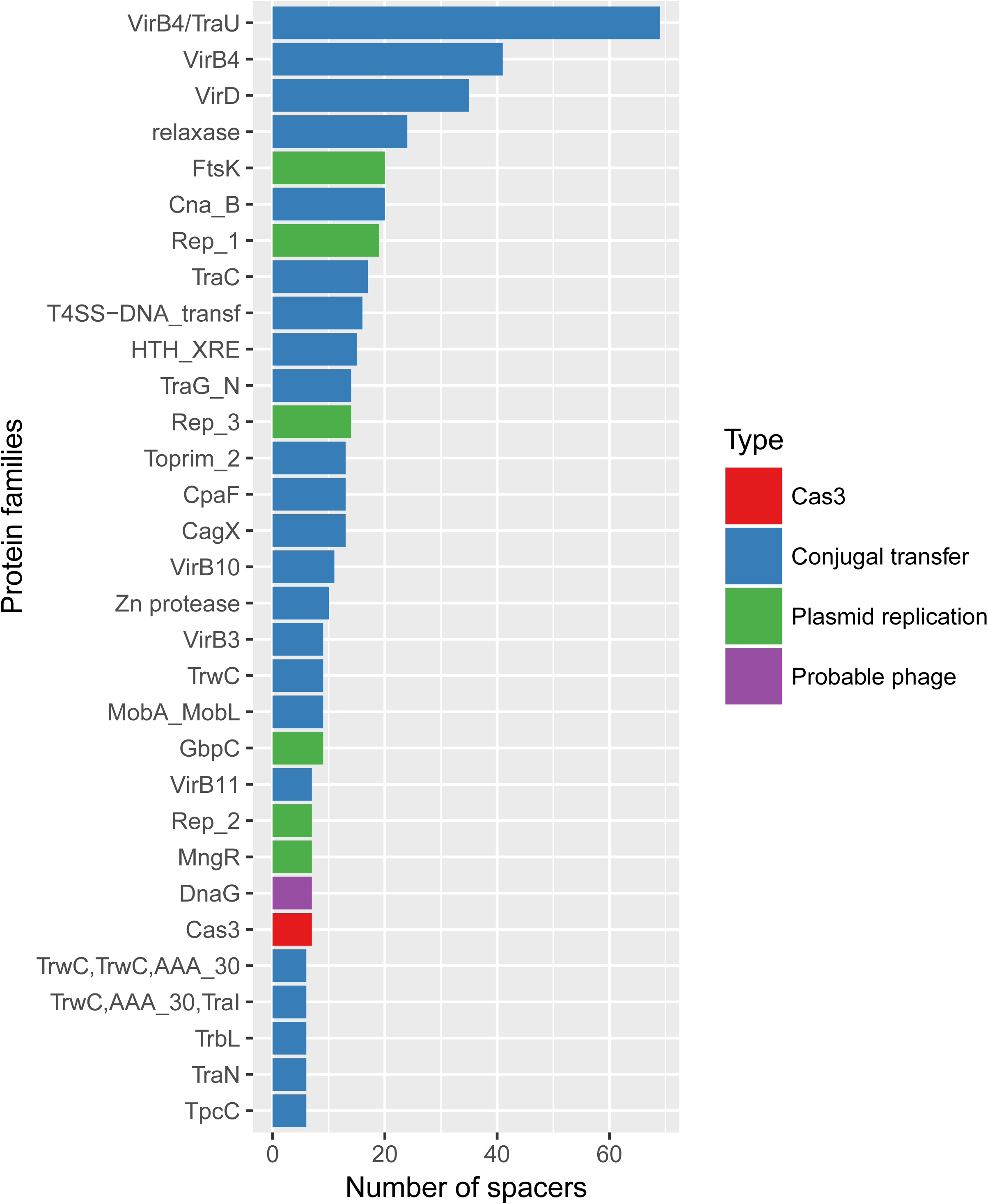
Breakdown of the protospacers from non-viral genes by gene family. Genes implicated in conjugal transfer of plasmids and plasmid replication, a putative phage gene (not annotated as such) and *cas3* gene are color-coded. The protein family names are from the CDD database.

Where do the ∼93% of the spacers that comprise the dark matter of CRISPR arrays come from? In an attempt to gain insight into the origin of these spacers, we compared the nucleotide compositions of the spacers, the respective prokaryotic genomes and the virus genomes containing the corresponding protospacers. The compositions of the three sequence sets showed near perfect correlation and were almost identical across the entire range of the GC-content; closely similar results were obtained regardless of whether all spacers or only spacers with matches were included (Figure 3A,B). Compatible results were obtained when we compared dinucleotide and tetranucleotide compositions among the same sequence sets using principal component analysis: all points formed a homogeneous cloud, without any detectable partitioning (Supplementary figures 4 and 5). Given the wide range of the GC-content covered, from ∼20 to ∼70% and the near indistinguishable features of the three sets of sequence, these observations strongly suggest that they all come from a single, intermixing, species-specific sequence pool. Bacteriophage genomes are generally considered to have a lower GC-content than the host genomes such that prophages form AT-rich genomic islands ^23^, which seems to be at odds with the near perfect correlation we observed. To investigate this discrepancy, we compared the GC-content of phage and host genomes for several bacteria for which numerous phages have been characterized; all available phage genomes were included in this analysis, regardless whether or not corresponding spacers were detected. In most cases, there was indeed considerable AT-bias in phages but numerous phage genomes had the same composition as the host and spacers (Figure 4). Conceivably, the spacers come from the most abundant phages that match the hosts in the GC-content.

**Figure 3.**
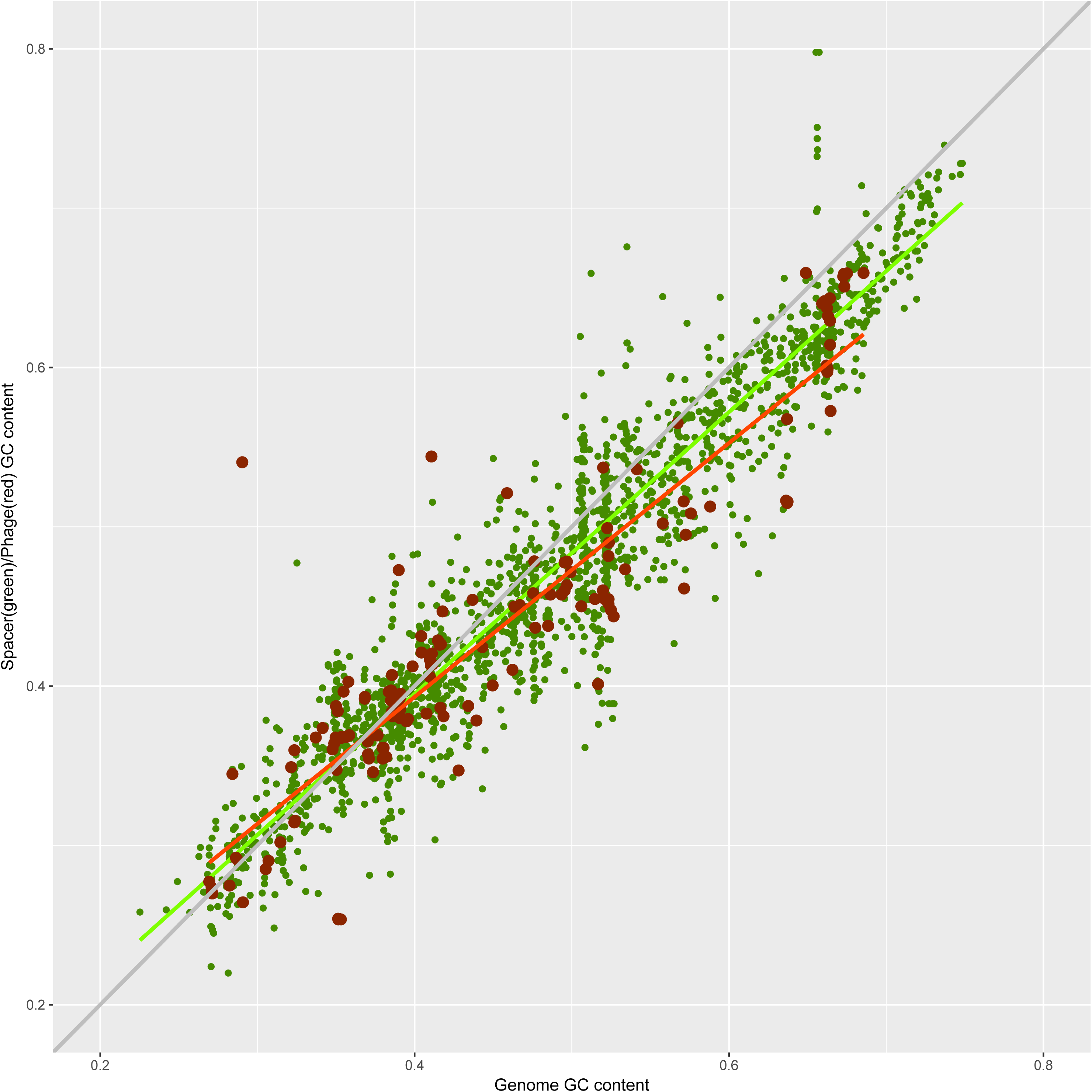
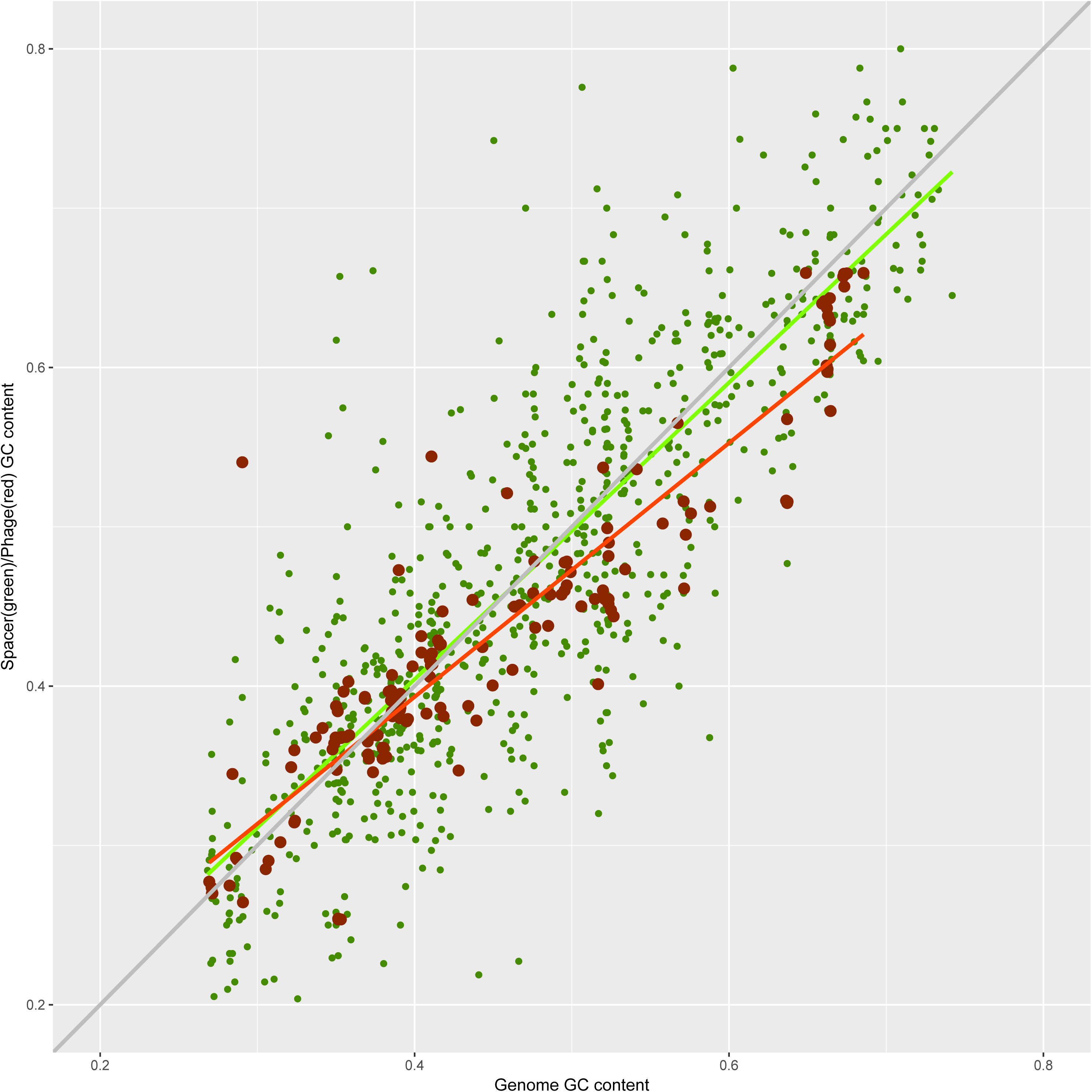
Correlations between the nucleotide compositions of spacers, the genomes of the respective microbes and their viruses. A. GC-content of spacers vs GC-content of microbial genomes and viruses B. GC-content of spacers with matches vs GC-content of microbial genomes and viruses Linear trend lines are shown for the GC-content of spacers (green) and viral genomes (red), and the x=y line is included to guide the eye.

**Figure 4.**
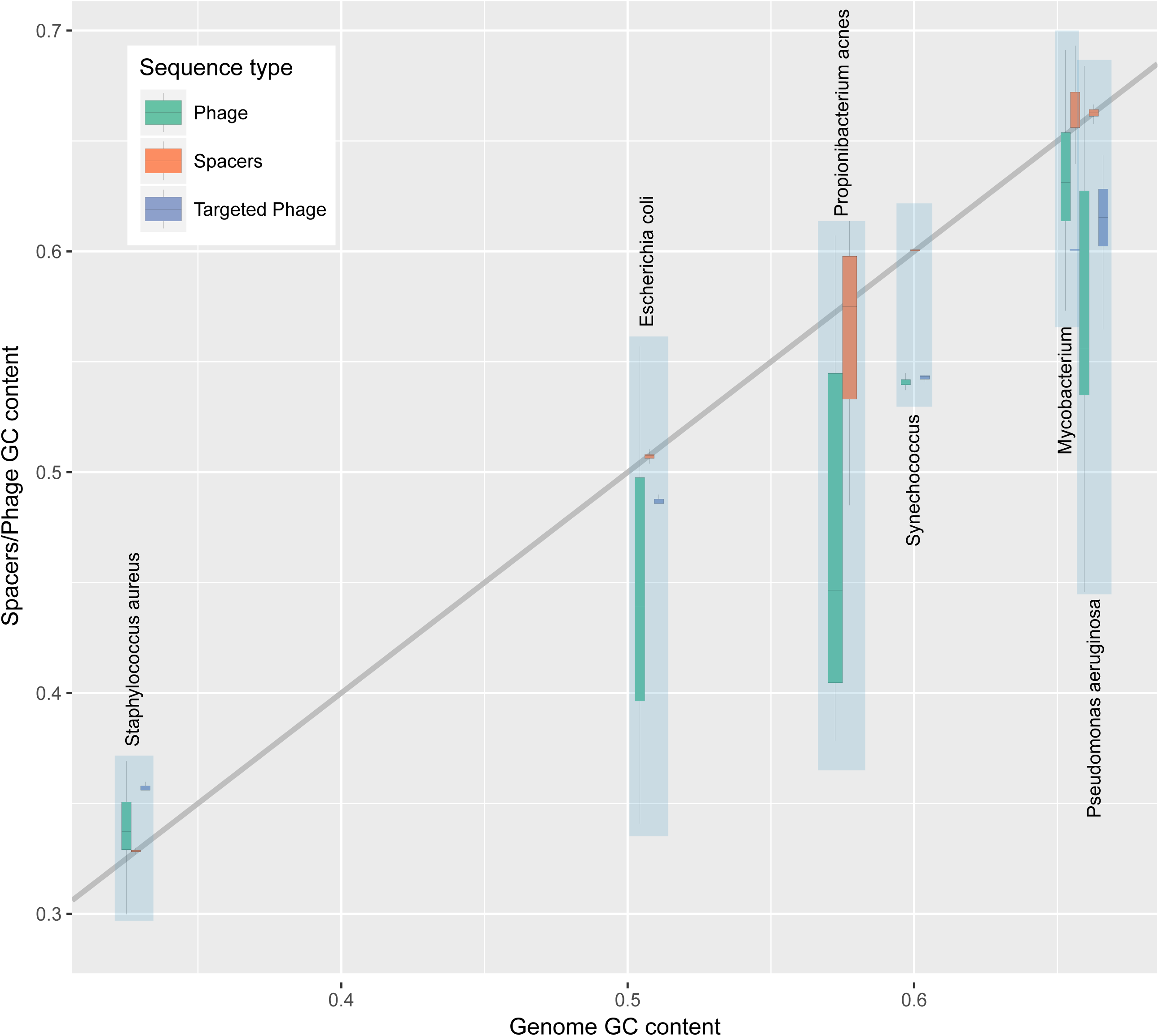
Correlations between the nucleotide compositions of spacers, genomes of bacteria with numerous characterized viruses and the corresponding viral genomes.

We further investigated the provenance of the dark matter spacers using an alternative approach. Matches to genomes from different microbial taxa, in the range from strains within the same species to different domains (archaea and bacteria), were tallied for the CRISPR spacers and for ‘mock spacers’, i.e. 1000 randomly sampled sequence segments of the same length from each CRISPR-carrying genome. The distributions of the matches were substantially different for the two sequence sets: the spacers matched genomic sequences almost exclusively within the same species, and almost none were found outside the same genus, whereas for the mock spacers, numerous matches were detected in distantly related genomes (Figure 5A). The distributions of the number of matches per (mock) spacer are quite different also, with the spacers being largely unique or matching only a few sequences, in contrast to the distribution for the ‘mock spacers’ that was dominated by a peak of abundant matches (Figure 5B). These observations indicate that the protospacers come from a sequence pool that is sharply different from the average genomic sequence in terms of evolutionary conservation. The protospacer sequences are extremely poorly conserved, which is the property of the mobilome.

**Figure 5.**
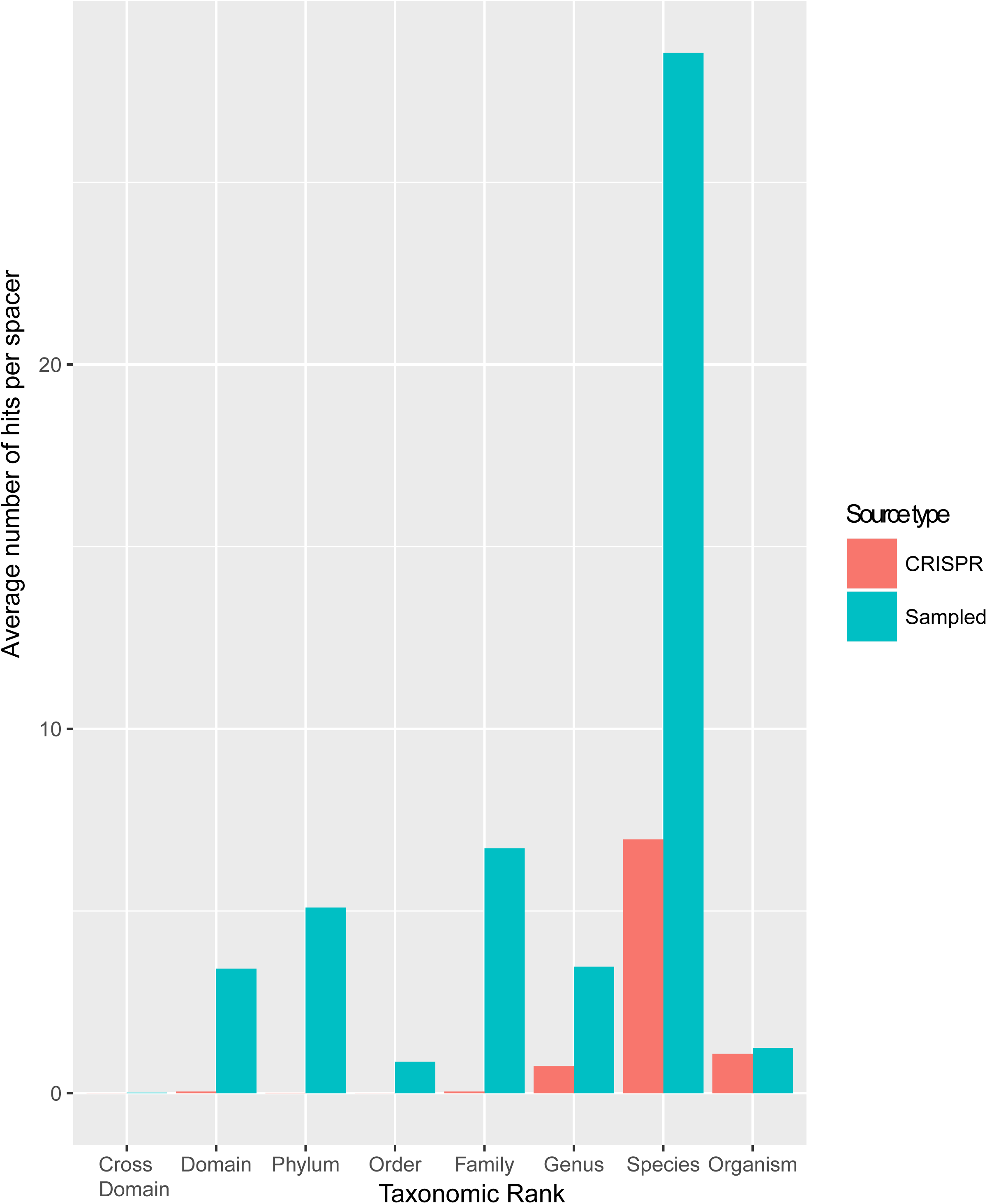
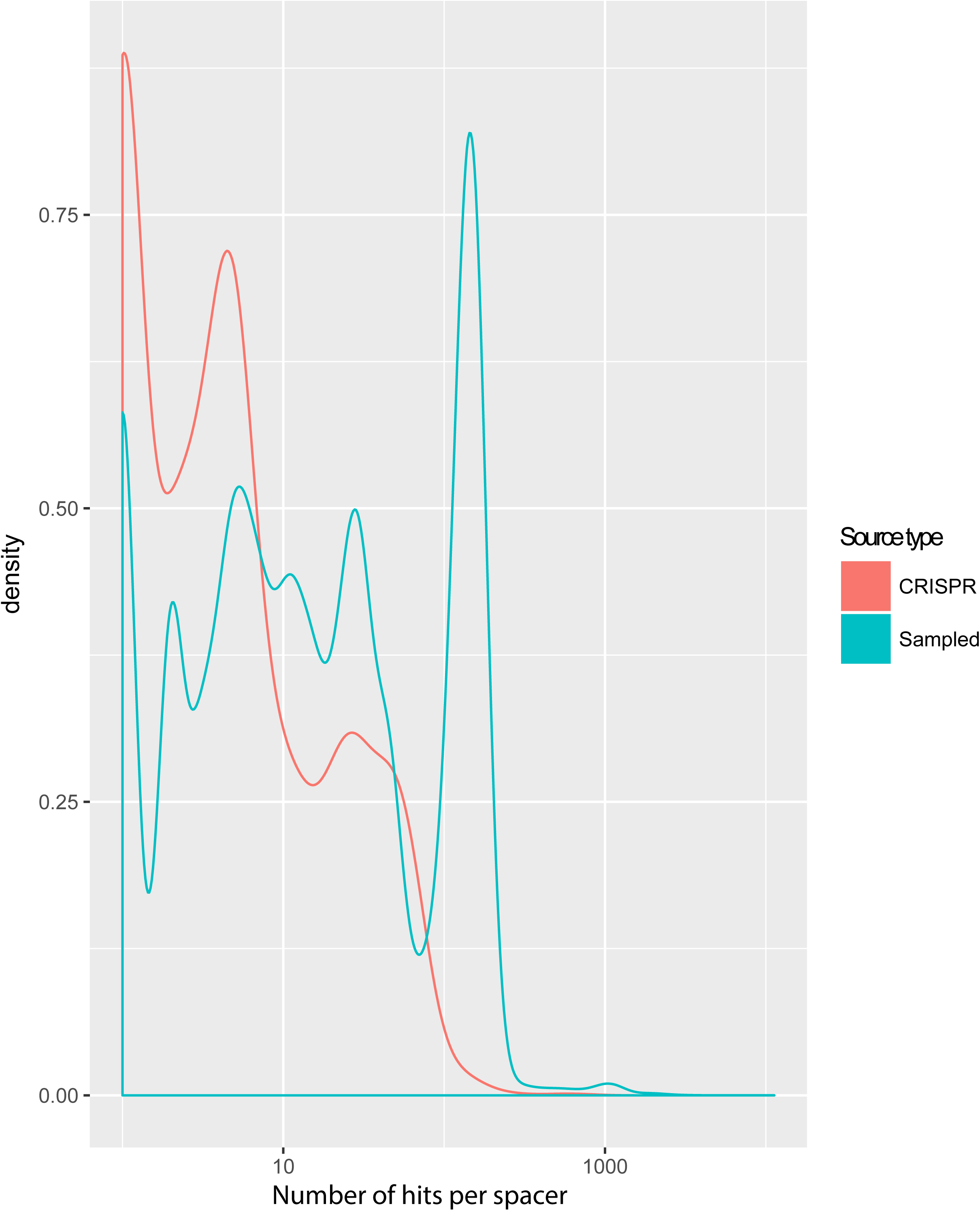
Spacer sequence conservation compared to the genomic average. A. Distribution of matches for the spacers and the ‘mock spacers’ across the microbial taxonomic ranks B. Distributions of the number of matches to the same species per spacer for the spacers and the *‘*mock spacers*’*

In the present dissection of the CRISPR (proto)spacer space, we made two principal observations. First, the spacers with detectable protospacer matches that persist in CRISPR arrays originate (almost) exclusively from genomes of mobile elements, mostly viruses, but also plasmids. This is not an unexpected finding, being compatible with multiple previous observations on individual prokaryotic genomes, but the overwhelming dominance of mobilome-derived sequences is now validated quantitatively on the scale of the entire prokaryotic sequence space. Notably, the great majority of viral protospacers were actually detected in provirus sequences. In part, this could reflect bias caused by the incompleteness of the current virus sequence database but the possibility also presents that CRISPR-Cas systems play a particularly important role in the control of provirus induction. Such a mechanism is suggested by the demonstration of transcription-dependent targeting of viral genomes by some CRISPR-Cas systems ^24^.

The strong selectivity of the CRISPR-Cas systems towards the mobilome is likely to stem from two sources, namely, self vs non-self discrimination at the stage of spacer incorporation and selection (preferential survival) of microbial clones incorporating non-self spacers. The mechanisms of discrimination remain far from being perfectly understood but at least some preference for non-self genomes through recognition by the adaptation complex of actively replicating and repaired and/or transcribed DNA has been demonstrated ^24^. Selection appears to be important as well because, when the nuclease activity of the effector is abolished, self-matching spacers accumulate ^25^. The relative contributions of self vs non-self discrimination and selection to the dominance of the mobilome as the source of detectable protospacers remain to be assessed and are likely to differ across the diversity of the CRISPR-Cas systems. Regardless, the result is a (near) complete exclusion of ‘regular’ microbial sequences from the spacer space. This exclusion involves not only the host but also other microbes, suggesting that CRISPR provide protection from viruses and on many occasions prevent plasmid spread but might not create a barrier for horizontal gene transfer via other routes, such as transformation.

The second key finding of this work is the demonstration that CRISPR spacers, both those with matches and the dark matter, the respective microbial genomes and their viruses belong to the same genomic pool as determined by (oligo-)nucleotide composition analysis. Together with the dominance of viral and plasmid sequences among the protospacers, these observations lead to the extrapolation that the overwhelming majority, and possibly, nearly all spacers originate from the same source, namely the species-specific mobilome. Then, whence the dark matter? There seem to be two complementary explanations. First, the dramatic excess of spacers without matches over those with detectable protospacers implies that for most microbes, the ‘pan-mobilome’ that they encounter in the course of evolution is vast and still largely untapped. Second, the lack of spacer matches could be caused by progressive amelioration of the spacer sequences caused, primarily, by mutational escape of viruses, which results in the loss of information that is required to recognize protospacers, at least in a database search. In the biological setting, spacers with mismatches can still be employed for interference and/or primed adaptation ^26^-^28^. Again, the relative contributions of the two factors remain to be investigated. The importance of amelioration is implied by the precipitous decline of the fraction of spacers with matches from the beginning towards the middle of arrays (Figure 1). Furthermore, in *Escherichia coli*, the only microbe, for which the virome can be considered comprehensively characterized, there are virtually no spacers with matches to the known viral genomes, suggesting that the apparently inactive CRISPR arrays in this bacterium have accumulated mismatches to the cognate protospacers that render them unrecognizable ^29^. Further characterization of the ‘pan-mobilomes’ of diverse bacteria and measurement of the spacer amelioration rates should improve our understanding of the evolution of the CRISPR spacer space and the virus-host arms race.

## Methods

### Prokaryotic Genome Database

Archaeal and bacterial genomic sequences were downloaded in March 2016 from the NCBI FTP site (ftp://ftp.ncbi.nlm.nih.gov/genomes/all/). The pre-computed ORF annotation was accepted for well annotated genomes (coding density >0.6 coding sequences per kilobase), and the rest of the genomes were annotated using Meta-GeneMark ^30^ with the standard model MetaGeneMark_v1.mod (Heuristic model for genetic code 11 and GC 30). The resulting database consisted of 4,961 completely assembled genomes and 43,599 partial, or 6,342,452 nucleotide sequences altogether (genome partitions, such as chromosomes and plasmids, and contigs).

### Detection and annotation of CRISPR arrays

All contigs from the prokaryotic genome database were analyzed with CRISPRFinder^31^ which identified 61,581 CRISPR arrays and PILER-CR ^32^ which identified 49,817 arrays. Arrays were merged by coordinates (CRISPRFinder array annotation was taken in case of overlap), which produced a set of 65,194 CRISPR arrays.

CRISPR-Cas types and subtypes were assigned to CRISPR arrays using previously described procedures 16,33. All ORFs within 10 kb upstream and downstream of an array were annotated using RPS-BLAST ^34^ with 30,953 protein profiles (from the COG, pfam, and cd collections) from the NCBI CDD database ^35^ and 217 custom CRISPR-Cas protein profiles ^33^. In cases of multiple CRISPR-Cas systems present in an examined locus, the annotation of the first detected variant was used to annotate the array.

Given the frequent misidentification of CRISPR arrays (Supplementary text 3), a filtering procedure for “orphan” CRISPR arrays (i.e. the arrays that are not associated with *cas* genes) was applied. A set of repeats from CRISPR arrays identified within typical CRISPR-*cas* loci was collected, and these were assumed to represent bona fide CRISPR (positive set). A BLASTN ^36^ search was performed for all repeats from orphan CRISPR arrays against the positive set, and BLAST hits were collected that showed at least 90% identity and 90% coverage with repeats from the positive set. All arrays that did not produce such hits against the positive set were discarded. The resulting 42,352 CRISPR arrays were used for further analysis.

### Detection of Protospacers

A set of unique spacers was extracted from the 42,352 CRISPR arrays by comparison of the direct and reverse complement sequences. The full complement of CRISPR arrays contained 720,391 spacers in total, with 363,460 unique spacers.

A BLASTN search with the following command line parameters: “-max_target_seqs 10000000 -dust no -word_size 8”; was performed for the unique spacer set against the virus part (NCBI taxid: 10239) of the NR/NT nucleotide collection ^37^ and against the prokaryotic database described above. The hits with at least 95% sequence identity to a spacer and at least 95% sequence coverage (i.e. allowing one or two mismatches) were accepted as protospacers. This threshold was defined from the results of a comparison of the number of spacer BLAST hits into prokaryotic and eukaryotic virus sequences (Supplementary Figure 6), where eukaryotic viruses served as a control dataset for false predictions. The threshold was set at the lowest false discovery rate of 0.06. As a result, 2,981 spacer matches were detected in viral sequences and 23,385 matches in prokaryotic sequences.

### Annotation of protospacers in prokaryotic genomes

To identify protospacers that belong to proviruses among the 23,385 spacer matches obtained in the prokaryotic genomic sequences, the following procedure was applied:

- All ORFs within 3 kb upstream and downstream of a spacer hit were collected
- A PSI-BLAST ^36^ search for all ORFs from these loci against the virus part of the NR database ^37^, with the following command line parameters: “-seg no -evalue 0.000001 -dbsize 20000000”, was performed
- A protospacer was classified as (pro)viral if it overlapped an ORF with a match in the viral part of NR database or if two or more ORFs with matches in the viral sequence set were identified within the neighborhood of the protospacer

Among the 23,385 spacer matches in prokaryotic genomes, 19,704 spacers targeted ORFs, of which 16,819 of were classified as (pro)viral. Among the 3,679 spacer targeting intergenic regions, 2,799 were classified as (pro)viral.

The results obtained with this classification procedure were compared to those obtained with PhiSpy ^38^, a commonly used prophage finder tool (default parameters) for the protospacer matches identified in the 4,961 completely assembled genomes. Of the 1,240 spacer matches in complete genomes, 999 hits were identified as (pro)virus-targeting by the *ad hoc* procedure described above. Using PhiSpy, 902 spacers were mapped to proviruses, of which 819 overlapped with the set of 999 viral matches detected by the ad hoc method, indicating high consistence of the predictions by the two approaches.

The distribution of protospacers across CRISPR-Cas types and subtypes was obtained from the unique spacer set. In cases when a unique spacer was identified in CRISPR arrays from different subtypes, only one instance was counted. The same procedure was applied to estimate the distribution of protospacers among the bacterial and archaeal phyla.

### Annotation of spacers matches in non-viral ORFs

The 2,885 ORFs that were targeted by spacers but not classified as viral proteins were annotated with 30,953 protein profiles (COGs, pfam, cd) from the NCBI CDD database and 217 custom CRISPR-Cas protein profiles using RPS-BLAST (using evalue 10e-4). Profile hits were obtained for 1,616 ORFs. The 1,269 ORFs with no identified profile hits were clustered using UCLUST ^39^, with the similarity threshold of 0.3. To assign ORFs to COG functional categories, the same procedure was performed against the COG proteins profiles only ^40^. The summary statistics for the functional categories was assembled using the COG table and is available at ftp://ftp.ncbi.nih.gov/pub/COG/COG2014/static/lists/homeCOGs.html

### Bipartite host-virus network analysis

The set of 2,981 spacer matches in the viral part of the NT/NR nucleotide collection was used to build a bipartite network with two types of nodes: CRISPR hosts and targeted viruses. All CRISPR hosts from the same genus were collapsed into a single node. Edges between network nodes were assigned when a protospacer matching a spacer in a given host was identified in in a virus. The network was visualized using the Cytoscape software ^41^.

### Nucleotide composition analysis of hosts, spacers and viruses

Nucleotide composition analysis was performed with the dataset of 2,104 complete genomes that contained CRISPR arrays. Frequencies of mono-, di-and tetranucleotides were calculated in genome sequences. The standard “prcomp” function from the R package was used for Standard Multidimensional Scaling.

Species with the most extensively sampled viromes were identified from the “/host” tag in RefSeq database for double-stranded DNA viruses:

| Host | Number of phages in RefSeq |
| --- | --- |
| Escherichia coli | 144 |
| Pseudomonas aeruginosa | 103 |
| Staphylococcus aureus | 77 |
| Propionibacterium acnes | 42 |
| Synechococcus sp. | 21 |
| Mycobacterium | 21 |

and analyzed separately, together with the associated viruses.

### Comparison of the distributions of spacer and random fragment matches in prokaryotic genomes

The comparison of the matches distribution for spacers and random fragments was performed on 2,104 complete genomes that contained CRISPR arrays. For each genome, 1000 random fragments, with the length equal to the median length of spacers in the given genome, were extracted. A BLASTN search against the prokaryotic database was performed for these fragments and for spacers, with following parameters: “-max_target_seqs 10000000 -dust no -word_size 8”. Exact matches were selected for further analysis.

## Acknowledgements

SS, KSM, YIW and EVK are funded intramural funds of the US Department of Health and Human Services (to National Library of Medicine).

